# The small protein MntS evolved from a signal peptide and acquired a novel function regulating manganese homeostasis in *Escherichia coli*

**DOI:** 10.1101/2023.06.02.543501

**Authors:** Zachary Wright, Mackenzie Seymour, Kalista Paszczak, Taylor Truttmann, Katherine Senn, Samuel Stilp, Nickolas Jansen, Magdalyn Gosz, Lindsay Goeden, Vivek Anantharaman, L. Aravind, Lauren S. Waters

## Abstract

Small proteins (< 50 amino acids) are emerging as ubiquitous and important regulators in organisms ranging from bacteria to humans, where they commonly bind to and regulate larger proteins during stress responses. However, fundamental aspects of small proteins, such as their molecular mechanism of action, downregulation after they are no longer needed, and their evolutionary provenance are poorly understood. Here we show that the MntS small protein involved in manganese (Mn) homeostasis binds and inhibits the MntP Mn transporter. Mn is crucial for bacterial survival in stressful environments, but is toxic in excess. Thus, Mn transport is tightly controlled at multiple levels to maintain optimal Mn levels. The small protein MntS adds a new level of regulation for Mn transporters, beyond the known transcriptional and post-transcriptional control. We also found that MntS binds to itself in the presence of Mn, providing a possible mechanism of downregulating MntS activity to terminate its inhibition of MntP Mn export. MntS is homologous to the signal peptide of SitA, the periplasmic metal-binding subunit of a Mn importer. Remarkably, the homologous signal peptide regions can substitute for MntS, demonstrating a functional relationship between MntS and these signal peptides. Conserved gene-neighborhoods support that MntS evolved from an ancestral SitA, acquiring a life of its own with a distinct function in Mn homeostasis.

**Significance:** This study demonstrates that the MntS small protein binds and inhibits the MntP Mn exporter, adding another layer to the complex regulation of Mn homeostasis. MntS also interacts with itself in cells with Mn, which could prevent it from regulating MntP. We propose that MntS and other small proteins might sense environmental signals and shut off their own regulation via binding to ligands (e.g., metals) or other proteins. We also provide evidence that MntS evolved from the signal peptide region of the Mn importer, SitA. Homologous SitA signal peptides can recapitulate MntS activities, showing that they have a second function beyond protein secretion. Overall, we establish that small proteins can emerge and develop novel functionalities from gene remnants.

## Introduction

Although initially overlooked, bacterial small proteins (< 50 amino acids) and eukaryotic microproteins (< 100 amino acids) are now recognized to be numerous and act in diverse cellular processes from respiration to apoptosis (1–6). The small size of these proteins imparts distinct genetic and biochemical properties, such as evolvability, a tendency to fold only in certain environments, and unusual subcellular trafficking, while also making them challenging to study (1-3,6-8). Although few small proteins have been functionally characterized to date, those that have been studied often bind larger proteins and modulate their function (1–5). In bacteria, several small proteins are known to regulate membrane proteins, including transporters, kinases, and proteases (7,8). Small proteins can enable or prevent protease degradation of transporters or directly modulate their transport activity, often via the binding of their transmembrane helices (1,7,8). Yet even for these well-studied examples, little is known about small protein structures and binding determinants, how their activities are turned off, or how genes encoding them emerge in genomes (1,2). To address these questions, we investigated MntS, a small protein with a well-defined physiological role in regulating manganese (Mn) homeostasis.

Mn helps catalyze diverse chemical reactions in all cells and is important for certain redox reactions and sugar-phosphate-using enzymes, such as specific kinases and phosphatases (9–11). Mn is also necessary for processes such as the innate immune response, cyclic dinucleotide signaling, phage defense systems, stress-related signal transduction, carbohydrate metabolism, and photosynthesis (10–20). Additionally, Mn contributes to the detoxification of reactive oxygen species (ROS) (20–22). Due to these essential roles, many pathogenic and symbiotic bacteria require Mn to survive in eukaryotic host tissues (22–25). Other bacteria require Mn for cellular differentiation and energy production (19,20,26,27). However, despite its beneficial functions, excess Mn can be toxic, likely by displacing other metals and rendering key proteins inactive (10,28–32). Therefore, cells must monitor and respond rapidly to changing Mn levels to maintain optimal intracellular Mn concentrations.

Bacterial Mn homeostasis systems consist of Mn transporters and Mn-binding regulators that control transporter expression (22,25,29). Bacteria harbor two main types of high-affinity Mn importers, MntH, a member of the NRAMP family, and the ABC cassette importer SitABCD/MntABC (29,33,34). Strains lacking Mn importers show sensitivity to ROS, metabolic limitations, and reduced virulence (25,35). At least five classes of dedicated Mn exporters are present across bacteria, loss of which causes sensitivity to high Mn via mismetallation of critical proteins and often also causes decreased virulence (10,29,30,36).

Some bacteria also have a small 42 amino acid protein, MntS, which impacts intracellular Mn levels (28,37). Overexpression of MntS in the presence of Mn causes a severe growth defect accompanied by high intracellular levels of Mn, low levels of Fe, reduced amounts of Fe-loaded heme, and decreased heme enzyme activity (28,37). However, the molecular mechanism of how MntS affects Mn homeostasis has remained unknown. Our findings show that MntS binds and inhibits the MntP Mn exporter and also address outstanding issues about small protein downregulation, evolutionary origins, and bifunctionality.

While small protein binding has been shown to control the functions of larger proteins, it is not known what signals cause small proteins to dissociate from their partners to terminate their regulatory action once conditions change (1). In addition to binding MntP, we found that MntS can interact with itself in cells with Mn independently of MntP. Oligomerization of MntS by self-binding could preclude interaction with MntP and shut off its regulation of the transporter. These Mn-regulated interactions would allow MntS to bind and block MntP when environmental Mn levels drop, and prevent MntS inhibition of MntP when Mn levels rise.

How small open reading frames such as MntS become expressed and develop functionality in cells has also been an open question. Small proteins could arise from duplication and degeneration of an existing gene or emerge *de novo* from previously non-coding sequences (1,5,38). We provide evidence that MntS evolved from an ancestral signal peptide (SP) region. We found that MntS is related to the SP of specific SitA proteins of the SitABCD Mn importer system. The SitA SP regions that share specific sequence features with MntS can confer MntS activities, while the SP sequences of distant SitA proteins cannot. We propose this SP region developed a new activity in Mn homeostasis, which allowed its gene to be maintained as a standalone open reading frame (ORF) after the rest of the *sitABCD* operon was lost from the chromosome of certain gammaproteobacteria.

It has been speculated whether cleaved SP regions can have a second function, in addition to directing the secretion of their appended cargo protein into the periplasm or endoplasmic reticulum (39,40). Our finding that MntS-related SitA SPs expressed as independent small proteins can bind and inhibit MntP raises the possibility that certain SPs with distinctive features can serve as modulators of larger proteins. These observations add to the growing body of evidence suggesting that, rather than being discarded and degraded, certain cleaved SPs can double as small regulatory proteins.

## Results

### MntS requires MntP to cause Mn sensitivity

The small protein MntS from the gammaproteobacterium *Escherichia coli* was originally identified by virtue of its regulation by Mn levels and the perturbation of Mn homeostasis upon MntS misregulation (37). We wanted to understand the mechanism of how excess MntS causes aberrantly high Mn levels and hypothesized that MntS regulates the levels or activity of another larger protein, similar to other small proteins (1,2). As overexpression of MntS or loss of MntP cause similar phenotypes (28), we sought to investigate whether MntS affects the activity or levels of the MntP Mn exporter. Since the MntS phenotypes require high Mn to manifest, we could not simply delete the *mntP* gene as *ΔmntP* cells are highly Mn sensitive and do not grow at the required Mn concentrations (28,37). To avoid this problem, we replaced the endogenous *mntP* coding sequence in the chromosome with a heterologous Mn exporter gene, *mneA* from *Vibrio fischeri*, which can rescue the *ΔmntP* Mn sensitivity growth defect in *E. coli* (41). The *ΔmntP::mneA* replacement strain expressed the MneA protein under the control of the native *mntP* promoter and endogenous riboswitch regulation (42). MneA expression enabled the *ΔmntP* strain to grow with up to 0.3 mM MnCl2 (Figure S1).

Using *ΔmntP::mneA* strains lacking the MntP protein, we asked whether overexpression of MntS could still cause Mn sensitivity in the absence of MntP. In the WT strain expressing MntP, high levels of MntS caused robust Mn sensitivity, but in the *ΔmntP::mneA* strain lacking MntP, MntS expression no longer conferred a Mn-dependent growth defect (Figure 1A). This result demonstrates that MntS requires MntP to cause the Mn sensitivity phenotype, suggesting MntS inhibits MntP function.

**Figure 1:**
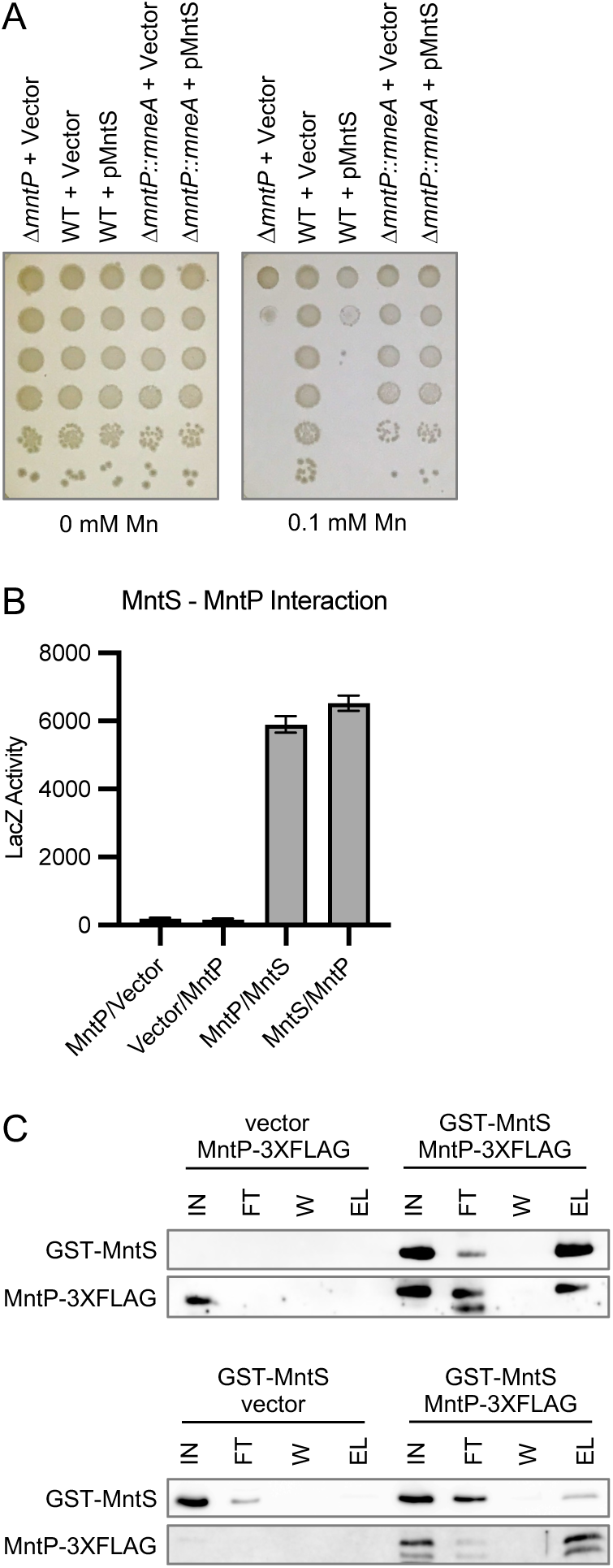
MntS requires MntP for Mn sensitivity and interacts with MntP *in vivo* and *in vitro*. **A)** Mn sensitivity assay showing that MntS can only cause a Mn-dependent growth defect in cells with the MntP protein. Overexpression of MntS causes a growth defect in the presence of Mn in the WT strain background, but the Mn sensitivity is lost in the Δ*mntP::mneA* strain which lacks the MntP exporter. Ten-fold serial dilutions of mid-exponential-phase cultures of WT, Δ*mntP* cells, or Δ*mntP::mneA* cells bearing the pBAD24 empty vector or pBAD24-MntS were spotted onto LB plates containing the indicated amounts of MnCl_2_. Results are representative of at least three independent replicates. **B)** Bacterial two-hybrid assay showing MntS interacts with MntP *in vivo*. Cells (Δ*cya* Δ*mntS*) containing two plasmids with fusions of MntS or MntP to the T18 and T25 fragments of adenylate cyclase were grown overnight in LB media. High β-galactosidase activity indicates efficient reconsistution of enzyme function and affinity between the fused pair of proteins. β-galactosidase assays were performed to quantify the LacZ activity representing the protein-protein interaction. Results are given in Miller units as the mean ± SD of three independent samples and are representative at least three independent experiments. **C)** Affinity co-purifications showing that MntS can pull-down MntP (top) and MntP can pull-down MntS (bottom). GST-MntS and MntP-3XFLAG were co-expressed in Δ*mntS* cells and purified on either glutathione sepharose or anti-FLAG resin. Proteins were separated by SDS-PAGE followed by immunoblotting using an anti-MntS antibody or anti-FLAG antibody.

### MntS directly binds to MntP

To examine whether MntS binds to MntP, we first used the *in vivo* adenylate cyclase bacterial two-hybrid system (43). We generated functional tagged MntP constructs with the T18 and T25 adenylate cyclase domains inserted into a cytoplasmic loop of the MntP protein. Point mutations in an acidic region of loop 3 of MntP have previously been shown to not disrupt its activity (41), and likewise MntP retained activity with insertions in this region (Figure S2). Two-hybrid assays with these MntP constructs and reciprocally tagged versions of MntS showed that MntS robustly interacts with MntP *in vivo* (Figure 1B). This interaction occurs with or without supplemental Mn.

We also found that MntS interacts with MntP using reciprocal affinity co-purifications with GST-MntS and MntP with 3XFLAG inserted into loop 3. Purification of GST-MntS also pulled down MntP-3XFLAG (Figure 1C, top), and similarly, purification of MntP-3XFLAG led to co-purification of GST-MntS (Figure 1C, bottom). Taken together with the observation that MntS acts through MntP (Figure 1A), these findings show that MntS can directly bind MntP and prevent its Mn export activity.

### MntS interacts with itself in the presence of Mn

While investigating the binding between MntS and MntP, we were surprised to find that MntS also interacted with itself (Figure 2A). Unlike the MntS-MntP interaction, the MntS self-binding required supplemental Mn at 0.3 mM or higher, and the interaction was specific to Mn. This result indicates that MntS can form a complex with itself under conditions of high intracellular Mn when its synthesis is normally repressed (37), suggesting self-oligomerization may be a method of shutting off its activity.

**Figure 2:**
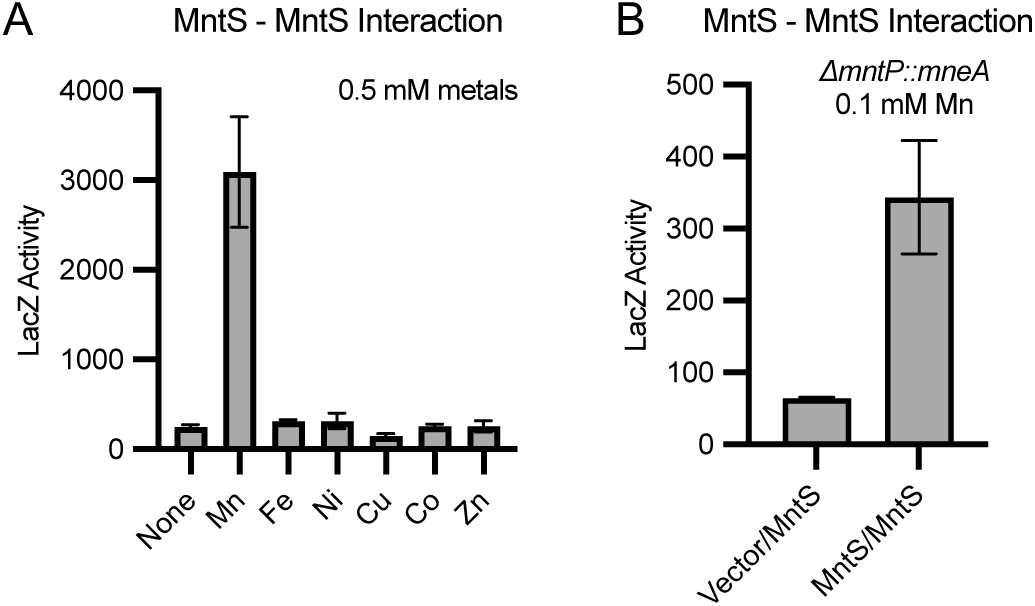
MntS oligomerizes in the presence of Mn independently of MntP. **A)** Bacterial two-hybrid assay showing MntS interacts with itself in a Mn-specific manner *in vivo*. Cells (Δ*cya* Δ*mntS*) containing two plasmids with fusions of MntS to the T18 and T25 fragments of adenylate cyclase were grown overnight in LB media with indicated amounts of metals. β-galactosidase assays were performed to quantify the LacZ activity representing the protein-protein interaction. Results are given in Miller units as the mean ± SD of three independent samples and are representative at least three independent experiments. **B)** Two-hybrid assay showing that MntS still interacts with itself in the *ΔmntP::mneA* strain which lacks the MntP exporter. Cells (Δ*cya* Δ*mntS* Δ*mntP::mneA*) containing two plasmids with fusions of MntS to the T18 and T25 fragments of adenylate cyclase were grown overnight in LB media with indicated amount of MnCl2. β-galactosidase assays were performed to quantify the LacZ activity representing the protein-protein interaction. Results are given in Miller units as the mean ± SEM of 7 or 16 independent samples (Vector/MntS or MntS/MntS) and are representative at least three independent experiments.

We also asked whether MntS could still bind to itself in the two-hybrid assay without MntP present. We observed that MntS could self-interact in a *ΔmntP::mneA* strain background (Figure 2B), consistent with an ability of MntS to directly bind to itself, possibly through its conserved hydrophobic residues, rather than oligomerizing on the surface of the MntP protein.

### MntS is homologous to the signal peptide region of certain SitA proteins

To determine conserved features of MntS for further functional studies, we used bioinformatic approaches to comprehensively identify MntS homologs. Interestingly, iterative sequence profile searches with *E. coli* MntS recovered not just MntS homologs, but also the SitA protein, a periplasmic subunit of a Mn importer. Initial searches found SitA proteins with significant similarity from other Enterobacterales, like *Rahnella* and *Sodalis*. Further transitive searches recovered additional partially related SitA proteins of other genera such as *Yersinia* (Figure S3A). The segment of similarity corresponded precisely to the signal peptide region of the SitA proteins (Figure 3A, S3A). However, not all SitA SPs shared unique sequence motifs with MntS. Other SitA SPs were more divergent and did not share the specific features with MntS (Figure 3A, S3A). Consistent with this, MntS was found within the phylogenetic clade containing the subset of SPs with which MntS shares specific motifs (Figure S3B).

**Figure 3:**
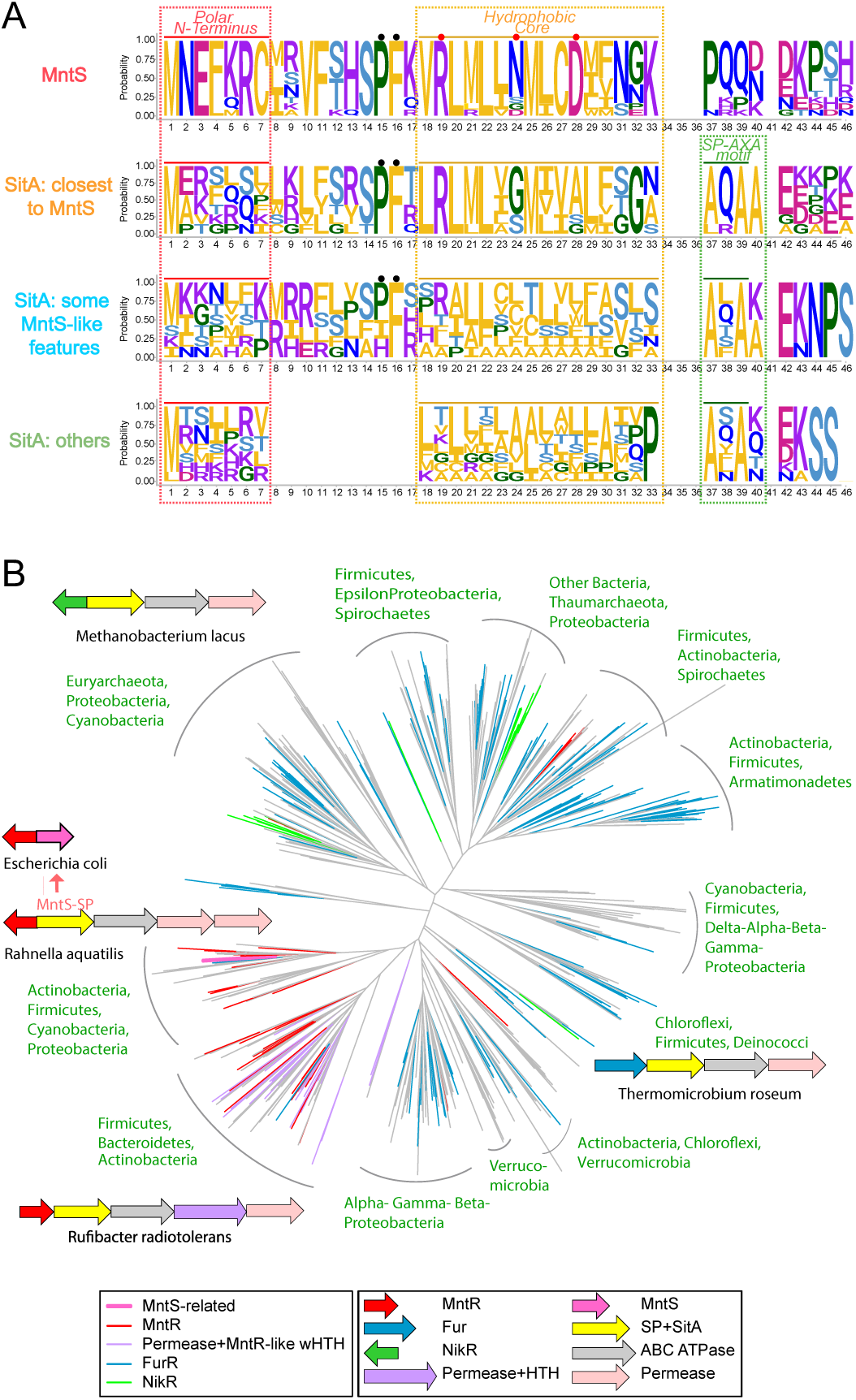
MntS is homologous to certain SP regions from SitA. **A)** A sequence logo displaying conservation of amino acid residues of MntS, the SitA SPs closest to MntS (e.g., *Rahnella* e=3 x 10^-10^, iteration 3; *Sodalis* e=5 x 10^-6^, iteration 4), SitA SPs with some MntS like features (e=10^-3^-10^-4^), and other SitAs with unrelated SPs. The letters represent amino acid abbreviations, with the height of each representing the probability of conservation in each of the indicated categories. Conserved amino acid positions, characteristic of each group, are shown with a dot above the residue: black dots mark the PF motif seen across MntS and related SPs, red dots indicate polar residues in MntS. The main features of a typical bacterial SP are boxed and labeled. **B)** A phylogenetic tree of SitA with representative operons, showing the associations of multiple distinct transcription factors with the *sit* operons and their presence across diverse bacterial clades. The 3-4 most frequent phyletic lineages of the major branches are shown in green; for complete phyletic patterns see Figure S4. The terminal branches are colored according to the transcription factors operonically linked to the SitA. Representatives of key operons are placed next to the clade of the tree where they are observed, with the organism shown below. Each arrow in the operon is a gene coding for the protein with the architecture indicated in the legend. The same colors shown in the legend are used for the corresponding genes (arrows) when shown in a representative operon. The *E. coli* MntS MntR dyad derived from the MntS-like SitA is also shown (pink arrow). Abbreviations: ABC-ATPase – ABC-ATPase subunit of the transporter;; NikR –TF with ribbon-helix-helix DNA-binding domain and C-terminal NikR-C metal-sensor; Fur – TF with N-terminal DNA-binding winged HTH (wHTH) domain and C-terminal Fur-C metal-sensing domain; MntR – TF with N-terminal wHTH domain and C-terminal DtxR-like metal-sensor domain. Permease+HTH – SitC/D like permease with MntR-like module fused to the cytoplasmic tail.

The SitA SPs related to MntS are similar in length to MntS, but notably both are longer than the other SitA SPs (40-45 vs. 25-34 residues). This length difference arises from the insertion of a 9-10 amino acid segment between the N-terminal polar region and the core hydrophobic helix (Figure 3A). Structural modeling indicates that MntS and the related SitA SPs likely adopt a bihelical structure with a kink at Pro15 at the end of the insert. Classical bacterial SPs show a tripartite organization with a N-terminal polar region on the cytoplasmic side of the membrane, a central hydrophobic region embedded within the membrane, and an AXA motif upstream of the signal peptidase cleavage site (Figure 3A). Although all SitA SPs, even those close to MntS, show these hallmark features, *bona fide* MntS has lost the final AXA motif and the downstream cleavage site. Further, a plot of the counts of hydrophobic vs polar residues clearly discriminates MntS from the SitA SPs (Figure S3C) and indicates a reduction of overall hydrophobicity in MntS relative to the SPs. Thus, we postulate that MntS emerged from a specific subset of SitA SPs via: (1) the loss of the secreted periplasmic binding domain; (2) degradation of the C-terminal AXA motif; (3) reduction of hydrophobicity of the central hydrophobic element which might have allowed its transition to being a soluble protein.

Intriguingly, SitA also functions in Mn homeostasis as the periplasmic binding protein of the ABC-cassette Mn importer system SitABCD (34). Given that the SP of secreted proteins is typically cleaved off during transport across the membrane by the signal peptidase and is then thought to be degraded (44,45), the region homologous to MntS would not be present in the mature Mn-binding SitA protein. Even the versions of SitA SPs closest to MntS show all the essential features of bacterial SPs, suggesting that they indeed function in the secretion of the mature SitA protein into the periplasm. However, the role of both MntS and SitA in Mn transport, and, notably, the MntS-like sequence features, structure, and length of this subset of SitA SPs raised the possibility that they might perform an additional MntS-like role.

### Genomic organization of Mn transporters and their regulators suggest the origin of MntS

To better understand the provenance of MntS, we conducted a comprehensive comparative genomics analysis of the *sitABCD* operon with *sitA* as the anchor. This operon is widely distributed across prokaryotes and is present in most major bacterial and certain archaeal lineages. The core of the operon encodes SitA, the secreted periplasmic metal-binding protein; SitB, the ABC ATPase pump driving import of the bound Mn; and one or two paralogs of the transmembrane permease subunits (SitC/SitD) (46). Phylogenetic analysis revealed that subclades of related SitA proteins are found in multiple distinct bacterial lineages (Figure 3B, S4). This, along with the presence of *sitABC/D* operons on plasmids, suggests that it has been widely disseminated through lateral transfer.

Additionally, about half of the SitABC/D operons are coupled in conserved gene neighborhoods with transcription factors (TFs) with distinct DNA-binding and metal sensor domains (Figure 3B, S4). Fur is the most common of these, seen in operons from across all major bacterial lineages (47). Its widespread coupling suggests that Fur was likely the ancestral TF of the SitABC/D systems. The next most prevalent coupling is with MntR (48). MntR appears to have displaced Fur on at least two occasions as the TF regulating the SitABC/D operon. The next prevalent association is with a NikR-like TF with MetJ/Arc (ribbon-helix-helix) DNA-binding domain (49), which has also displaced Fur on at least two occasions. A more limited association is with a metal-sensing AdcR-like TF displaying a MarR-like winged HTH DNA-binding domain (50). We also found more sporadic coupling to another TF related to LacI that might function as another alternative metal sensor. This pattern of TF displacement is consistent with the idea that coupling an operon with several alternative TFs brings genes under distinct regulatory regimes that help tailor their expression to inputs specific to an organism’s ecology (51).

One major subclade of SitA is primarily associated with MntR rather than Fur, and the SitA proteins with MntS-like SPs are all from within this subclade (Figure 3B). This observation indicates that MntS inherited its association with MntR from the ancestral SitA that was also associated with MntR, strengthening the evidence that MntS descended from the fragmented signal peptide of SitA. Importantly, all bacteria with complete genomes that have a standalone MntS or SitA SPs related to MntS also possess the target protein MntP, consistent with its second role in regulating MntP.

### MntS-like SitA signal peptides confer MntS phenotypes

To investigate if the SitA SPs related to MntS have an additional function, we cloned two such SPs (Figure S3A) and tested their ability to recapitulate MntS phenotypes in *E. coli*. First, we examined whether overexpression of the SitA SP regions could cause a growth defect in the presence of Mn, as is known for MntS (28,37). Interestingly, the SP from *Rahnella* SitA as well as the SP from *Sodalis* SitA conferred Mn sensitivity to cells (Figure 4A). As for MntS, the SP regions only caused a growth defect with Mn and not with other metals tested (Figure S5A). To investigate whether this was a general property of SP regions from SitA, we also examined the SP regions from *sitA* genes of *Salmonella*, *Shigella*, and *Klebsiella* which are distant from MntS (Figure S3A). These dissimilar SP regions did not cause Mn sensitivity (Figure 4A). Only the SitA SPs related to MntS were able to confer the growth defect in the presence of Mn, suggesting that the phenotype results from their specific sequence and structure similarity rather than general properties of SPs.

**Figure 4:**
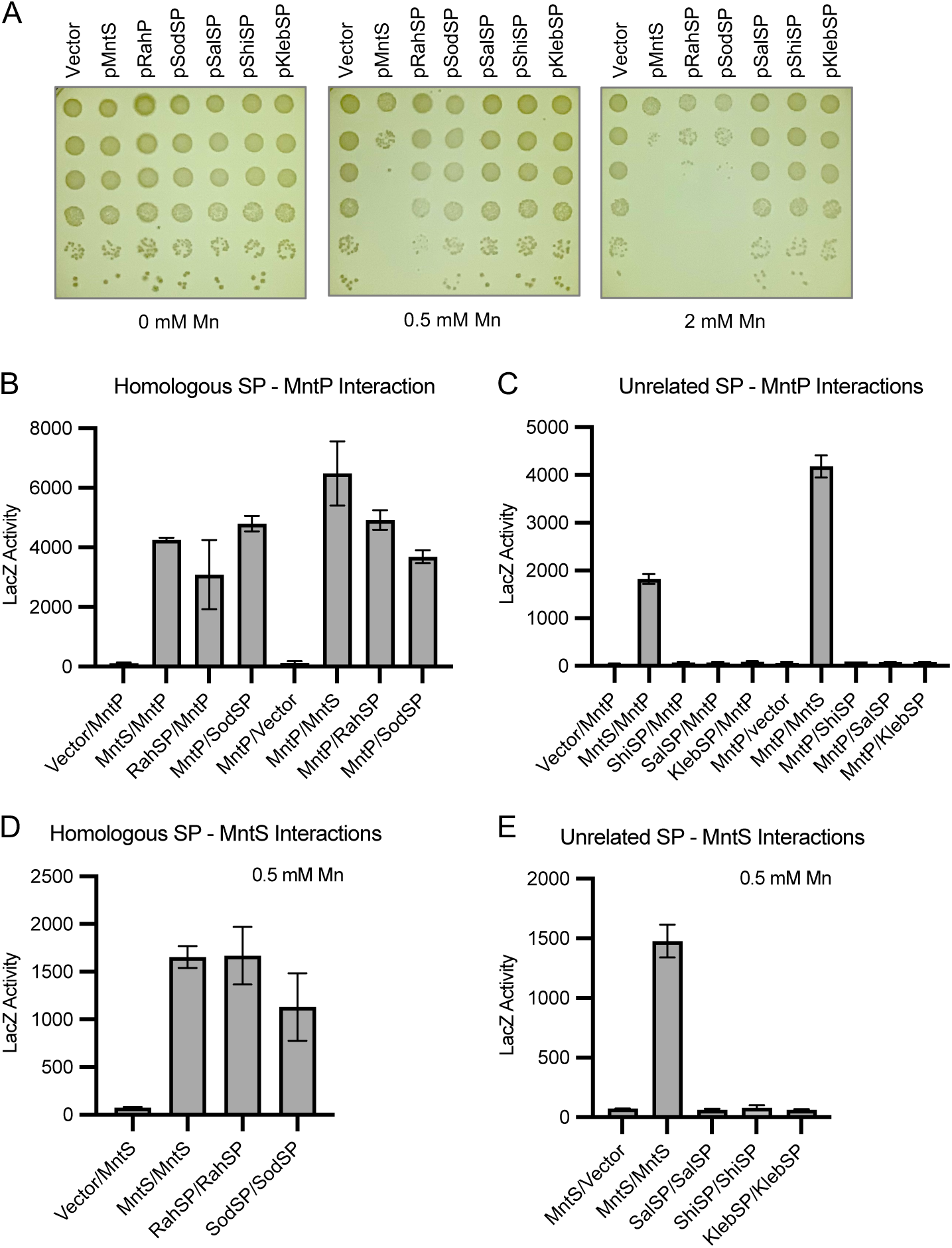
MntS-like SP regions of SitA recapitulate MntS activities. **A)** Mn sensitivity assay showing that overexpression of MntS or homologous SP regions from SitA proteins (*Rahnella* and *Sodalis*) can cause a Mn-dependent growth defect, while overexpression of unrelated SP regions from SitA proteins (*Salmonella*, *Shigella*, and *Klebsiella*) cannot cause Mn sensitivity. Ten-fold serial dilutions of mid-exponential-phase cultures of Δ*mntS* cells bearing the pBAD24 empty vector or pBAD24 expressing MntS or the RahSP, SodSP, SalSP, ShiSP, or KlebSP regions were spotted onto LB plates containing the indicated amounts of MnCl_2_. Results are representative of at least three independent replicates. **B)** Two-hybrid assay showing MntS and homologous SP regions from SitA proteins (*Rahnella* and *Sodalis*) can interact with MntP *in vivo*. For all two-hybrid assays (B-E), Δ*cya* Δ*mntS* cells containing two plasmids with fusions of MntS, the indicated SP regions, or MntP to the T18 and T25 fragments of adenylate cyclase were grown overnight in LB media with 0.5 mM MnCl2. β-galactosidase assays were performed to quantify the LacZ activity representing the protein-protein interaction. Results are given in Miller units as the mean ± SD of three independent samples and are representative at least three independent experiments. **C)** Two-hybrid assays showing unrelated SP regions from SitA proteins (*Salmonella*, *Shigella*, and *Klebsiella*) do not interact with MntP *in vivo*. **D)** Two hybrid assay showing that MntS and homologous SP regions from SitA proteins (*Rahnella* and *Sodalis*) can self-interact in cells with 0.5 mM MnCl2. **E)** Two-hybrid assay showing unrelated SP regions from SitA proteins (*Salmonella*, *Shigella*, and *Klebsiella*) cannot self-interact in cells with 0.5 mM MnCl2.

We next investigated whether the homologous SP regions acted through the MntP Mn exporter to affect cells, as observed for MntS (Figure 1A). Like MntS, the MntS-like SP region of *Rahnella* SitA also required MntP to cause Mn sensitivity (Figure S5B). Importantly, the MntP interaction with MntS was also recapitulated with the *Rahnella* and *Sodalis* SP regions (Figure 4B), but not with dissimilar SP regions from *Shigella*, *Salmonella*, or *Klebsiella* (Figure 4C). These data demonstrate that, like MntS, the homologous SP regions of SitA require MntP to cause the Mn sensitivity phenotype, bind MntP, and likely block its Mn exporter activity.

We further examined whether SitA SPs could dimerize in the two-hybrid assay, as we had demonstrated for MntS (Figure 2A). The MntS-like SPs from *Rahnella* and *Sodalis* displayed self-binding (Figure 4D), while the SPs from *Salmonella, Shigella,* and *Klebsiella* which are distant in sequence from MntS did not interact with themselves (Figure 4E). Like MntS, the *Rahnella* SP self-interaction required Mn and was Mn-specific (Figure S5C). Additionally, like MntS, the *Rahnella* SP, could bind to itself in a Δ*mntP::mneA* strain background (Figure S5D). Interestingly, MntS and the MntS-like SPs could interact with each other in all combinations (Figure S5E). Taken together, the ability of the SP regions of certain SitA proteins to affect bacterial growth in presence of Mn, to interact with and inhibit MntP, and to self-interact *in vivo* in the presence of Mn confirms that these SP sequences, when present as independent small proteins, can act similarly to MntS.

### Role of intact SP regions in the context of native full-length SitA protein

The fact that the N-terminal 42 amino acids corresponding to the SitA SPs function similarly to MntS when expressed as independent proteins suggests that the full-length SitA protein may also have this activity. To examine whether the *Rahnella* SP region still has MntS functionality when expressed from the native *sitA* gene, we expressed the full-length *Rahnella* SitA protein in *E. coli* (Figure 5A). We observed that the *Rahnella* SP region can also confer Mn sensitivity in the context of the full-length *Rahnella* SitA protein. As expected, the full-length *Salmonella* SitA protein, which has an SP distant in sequence from MntS, does not cause Mn sensitivity (Figure 5A). Chimeric constructs fusing the *Rahnella* SP region with the mature *Salmonella* SitA sequence starting after the signal peptidase cleavage site and vice versa revealed that only proteins containing the *Rahnella* SP caused Mn sensitivity (Figure 5B). Remarkably, MntS fused to SitA also was able to confer Mn sensitivity. Overall, these data strongly support the idea that MntS evolved from an ancestral SitA signal peptide region (Figure 6A). It also suggests that SPs of membrane and secreted proteins can possess additional regulatory functions.

**Figure 5:**
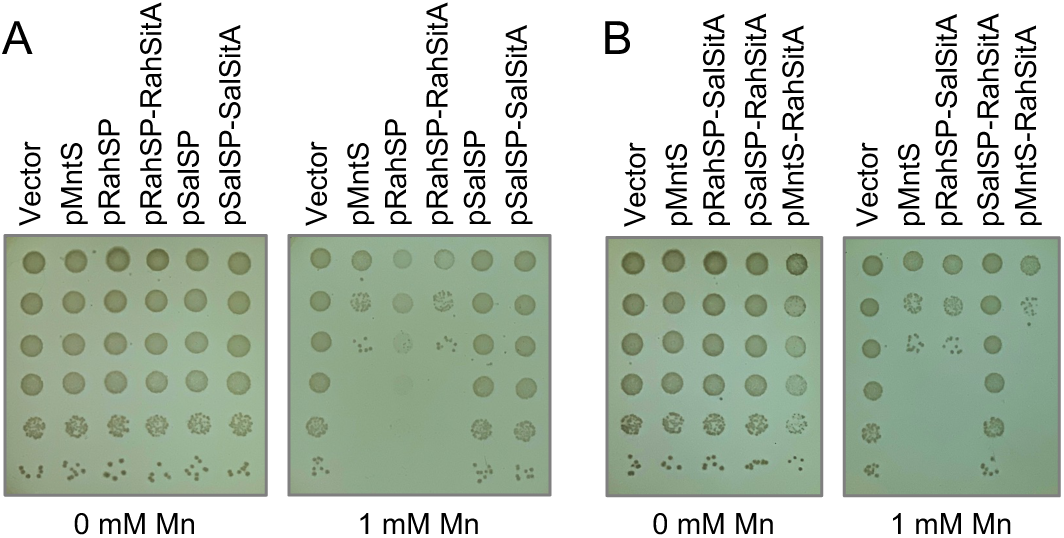
MntS-like SP regions of SitA confer Mn sensitivity as independent proteins or as part of the larger SitA protein. **A, B)** Mn sensitivity assay showing that MntS or the *Rahnella* SP region can cause a Mn-dependent growth defect when expressed as independent small proteins or as the N-terminus of the larger SitA protein. Ten-fold serial dilutions of mid-exponential-phase cultures of Δ*mntS* cells bearing the pBAD24 empty vector or pBAD24 expressing MntS, RahSP, SalSP, full-length Rah-SitA (SPRah-SitARah), full-length Sal-SitA (SPSal-SitASal), chimeric SPRah-SitASal, chimeric SPSal-SitARah, or chimeric MntS-SitARah were spotted onto LB plates containing the indicated amounts of MnCl2. Results are representative of at least three independent replicates.

**Figure 6:**
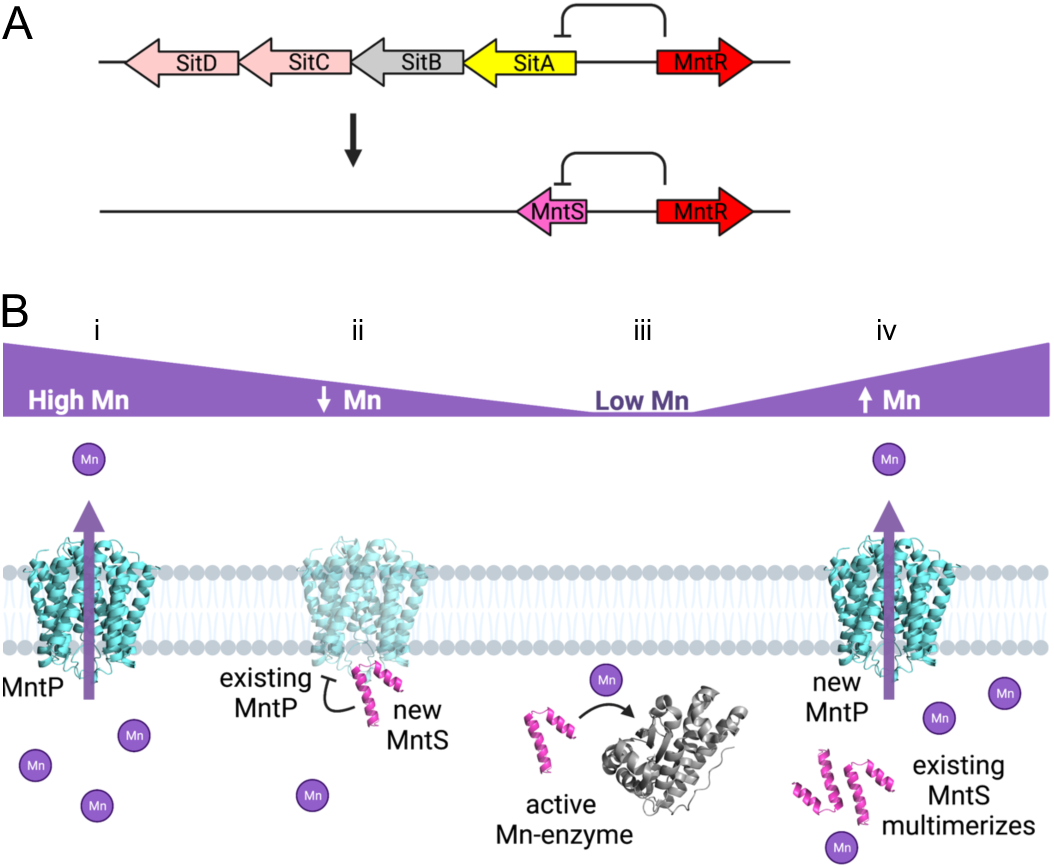
Model for MntS evolution and activity under different Mn conditions. **A)** Model for MntS origin from the SitA SP showing that in an ancestral gammaproteobacterial organism, most of the coding sequence for the *sitA* operon regulated by the MntR Mn-binding transcription factor was lost, leaving behind the N-terminal *sitA* region. This SP sequence was still regulated by Mn levels and evolved into the independent small protein MntS. **B)** Model for MntS activity and protein-protein interactions in different Mn conditions. **i)** Under steady-state high Mn levels, MntP is synthesized and exports excess Mn out of the cell. MntS is not present, as its expression is repressed by Mn-MntR (37). **ii)** As cells move from high Mn to low Mn, MntP expression is no longer induced and MntS begins to be synthesized. During this transition, MntP and MntS would be simultaneously present in cells. Although MntP synthesis is shut off, existing MntP would still be present in cells, which would be a liability since Mn levels are no longer in excess and the cell would need to retain some intracellular Mn for future starvation levels. As part of a Mn sparing response, MntS would bind and inhibit the existing MntP. **iii)** Under steady-state low Mn levels, MntP is not present but MntS is expressed to aid known Mn-using enzymes (Mn-SOD and Mn-RNR) to acquire their cofactor to be functional during Mn starvation (28). **iv)** As cells move from low Mn to high Mn, MntP begins to be synthesized and MntS expression is once again repressed. During this transition, MntP and MntS would be simultaneously present in cells. As MntP activity is needed to counteract the toxic Mn conditions, MntS should be prevented from inhibiting MntP. Under these increasing Mn conditions, MntS self-interacts (Figure 2A). We propose with high Mn, the MntS-MntS homodimer is more stable than the MntS-MntP heterodimer. This self-interaction may inactivate the MntS protein and prevent it from blocking MntP export of Mn.

## Discussion

### Extensive regulation of MntS and MntP activity during dynamic Mn fluctuations

Since Mn is both required for cells and toxic in excess, its levels in cells must be carefully and dynamically controlled. Mn homeostasis and intracellular availability are regulated at many levels, including well-studied transcriptional control of Mn transporters as well as newly recognized roles for RNA-binding proteins, small RNAs, and protein stability of transporters (30). Here we now report the role of a small protein, MntS, in regulating a Mn exporter. Previously, inappropriate expression of MntS was shown to perturb intracellular Mn and Fe pools via an unknown mechanism, which led to poor growth in Mn and loss of heme-enzyme activity (28). This study demonstrates that MntS Mn sensitivity only occurs in the presence of MntP and that MntS binds to MntP to downregulate its activity (Figure 1). The finding that Mn efflux is regulated by a small protein emphasizes that controlling Mn levels through export is a high priority for bacteria, as the MntP protein is also transcriptionally regulated by MntR and post-transcriptionally by the *yybP-ykoY* riboswitch (37,42).

Uniquely we also suggest a mechanism for how MntS shuts off its own activity. While small protein expression is commonly induced by environmental signals, it is not clear how small protein activity is downregulated once the cellular stress has been relieved (1). To release the target protein from regulation, small proteins must disassociate from the larger protein and/or be degraded. Since regulators should have dynamic protein-protein interactions with their targets, one way to disrupt the small protein binding to its target protein could be titrating away the small protein via another binding event when environmental conditions change. In addition to binding the MntP transporter, the MntS small protein also binds to itself in a Mn-dependent manner in cells (Figure 2A). We suggest that this self-binding causes MntS to release MntP when Mn levels rise. It is tempting to speculate that Mn binds MntS, which enables it to oligomerize either by Mn-induced structural changes or Mn bridging between monomers. Alternatively, Mn may stabilize MntS by preventing its degradation, allowing its levels to accumulate sufficiently to self-interact. Whatever the mechanism, we propose the MntS-MntS complex is mutually exclusive with a MntS-MntP interaction, coupling an environmental signal (high Mn) to the inhibition of small protein regulation via structural changes.

Although we have shown that MntS binds MntP and inhibits its activity, both proteins would not typically be present in the cell at the same time under steady-state conditions. MntS expression is repressed by high intracellular Mn levels, while MntP is induced by high Mn (28,42). However, during transitions from high to low or low to high Mn levels, both proteins would transiently exist in a cell, allowing for MntS to interact with MntP and alter its activity (Figure 6B). When cells move from a Mn-replete to a Mn-deficient environment, MntP synthesis would be halted by the absence of transcriptional induction via Mn-loaded MntR and lack of translational activation via the Mn-bound *yybP-ykoY* riboswitch (37,42). However, existing MntP would pose a threat by removing remaining Mn from cells that will shortly be starved for Mn. The remaining MntP could be bound by newly synthesized MntS to inhibit its Mn export activity, retaining some intracellular Mn.

Conversely, when cells transition from low Mn availability into excess Mn conditions, MntS synthesis would be shut off by Mn-MntR transcriptional repression (37). MntP expression would be induced by Mn-MntR and the *yybP-ykoY* riboswitch to enable cells to remove excess intracellular Mn. However, existing MntS could still inhibit MntP activity, which would be detrimental to bacteria needing to avoid Mn toxicity. The observation that MntS oligomerizes in the presence of high external Mn (Figure 2A) may provide an explanation for how MntS inhibition of MntP is avoided in this circumstance. Binding of MntS to itself could be mutually exclusive with binding to MntP, causing MntS oligomerization to shut off a potential inhibition of MntP (Figure 6B).

### Relevance of MntS for the models of the origin of small proteins

We show that MntS emerged by fragmentation of the larger SitA protein and acquired a role of its own as a small protein (Figure 6A). It is notable that SitA is a Mn-binding subunit of a Mn importer, while MntS is an inhibitor of the Mn exporter, MntP. We postulate that the signal peptide of certain SitA proteins in gammaproteobacteria acquired a secondary function to counter Mn efflux so that there could be a smooth shift to influx in Mn-requiring conditions (Figure 6). Our results suggest that MntS emerged via a bifunctional intermediate that initially played the role of both a SP (ancestral function) and a regulator that was cleaved off from a larger precursor (neomorphic function). Similarly, the small protein MgtR, which negatively regulates the stability of the MgtC magnesium transporter, also appears to have originated from a breakaway signal peptide [(52),(LA unpublished observations)]. Thus, this model for the origin of small proteins through multifunctionality and subsequent fragmentation could be a more general phenomenon.

The presence of regular repeating units within proteins, such as α-α bihelical hairpins and metal-stabilized elements, indicates that widespread ancestral protein domains arose from the duplication and accretion of small primordial proteins (53,54). These progenitor proteins are inferred to have comparable size, structure, and stabilization determinants (e.g., metal chelation) as extant small proteins. However, barring a few ribosomal small proteins, none of the extant small proteins are found broadly across the tree of life, which indicates they have arisen more recently (55). This suggests the processes that assembled the first protein domains from primordial small proteins could also work in reverse, through fragmentation of existing proteins to continually produce new small proteins, as exemplified by MntS.

Small protein evolution via fragmentation of larger proteins has advantages over acquisition of neomorphic function by *de novo* ORFs. The small protein can inherit the expression mechanisms and functional “pre-adaptations” developed by the parent protein. Moreover, fragments of larger proteins are more likely to fold into a functional structure than the product of *de novo* ORFs (56). The decreased structural stability and increased surface hydrophobicity of small proteins could drive protein-protein interactions, including multimerization or metal binding, to acquire a more stable state (57) (58,59). Consistent with this, a small protein regulating aconitase was shown to emerge from its protein-protein interaction region via duplication and fragmentation of the ancestral aconitase gene (57).

Our gene neighborhood approach could also provide a way to assign a cognate metal to divergent versions of metal-binding proteins. In many bacterial species, the physiologically relevant metal-binding specificity of the SitA-like solute binding protein component of ABC transporters is not clear (22,60,61). Indeed, identifying the physiologically relevant metal for metal-binding proteins like transporters from their sequence or structure alone is a notorious challenge (22,31). Our findings that the *sitABC/D* operon is associated with different metal-sensing transcription factors across bacterial species (Figure 3B, S4) suggests that using large-scale genomic organization of transporters could be a useful way to discern the metal specificity of a distinct protein sequence in its particular organism. In addition, novel modes of regulation can be uncovered by comparative genomics. For example, we found that a MntR-like module directly fused to the cytoplasmic tail of the permease subunit SitC/D in certain operons (Figure 3B, S4). These versions might provide an intracellular allosteric metal-binding site that could regulate import in addition to a potential transcription role. Similarly, a subset of the *sitABC/D* operons encode a further two-gene module coding for a HypB/CobW/UreG-like GTPase (46,62) and a WD40 β-propeller protein (Figure S4). These are likely together constitute a chaperone for the insertion of the metal imported via the transporter system into the metallocenters of enzymes.

### Signal peptide regions with second functions

After observing the similarity between MntS and certain SitA SPs, we showed that MntS-like SP regions of SitA can bind to and inhibit the MntP Mn exporter protein. Moreover, the SP region is functional when expressed either as an independent small protein or as the N-terminus of the full-length SitA protein (Figure 5). This suggests the SP regions of these SitA proteins have a second function, beyond directing the secretion of the mature SitA protein into the periplasm. Further, our findings indicate that SP regions of other periplasmic binding proteins could also have regulatory functions as independent small proteins after being cleaved off from their parent proteins. It is not clear how and where such cleaved SP regions would accumulate in cells to carry out their functions. While separated SP regions are generally assumed to be degraded by signal peptide hydrolases in bacteria (e.g., SppA and RseP), this has primarily been studied for model SP substrates and has not been broadly examined (45,63,64). Intramembrane cleavage may also be a processing step that generates a SP with an additional functional regulatory role. The SppA protease processes lipoprotein signal peptides to generate extracytoplasmic quorum sensing signaling molecules and RseP cleaves single-pass TM proteins to liberate cytoplasmic forms that control stress responses, including iron homeostasis (40,65). A few viral and eukaryotic SP regions are also known to have additional functions as signaling molecules once processed and released from the membrane (39,40). Overall, the extent to which SP regions carry out alternative regulatory roles once cleaved off their cargo protein, as well as how these SP regions escape degradation are open and intriguing questions (39,40).

## Materials and Methods

### Strains and Plasmids

Strains, plasmids, and oligonucleotide primers used in this study are listed in Table S1, S2, and S3. *E. coli* cultures were routinely grown in Luria broth (LB) with 150 μg/mL ampicillin, 30 μg/mL kanamycin, 30 μg/mL chloramphenicol, 0.5 mM IPTG, or 0.2% L-arabinose as needed. Chromosomal deletion strains were generated using pKD4 or pKP1 via mini-λ Red recombineering (66,67). Deletion constructs were moved into other genetic backgrounds by P1 phage transduction. When needed, the kanamycin resistance marker was removed via the FRT sites with pCP20 (68).

Plasmids were generated by cloning PCR amplified genes into pBAD24, pKT25c, or pUT18c (43). Sequences coding for *mntP, mneA,* or *sitA* were amplified by PCR from bacterial chromosomal DNA or from synthetic DNA fragments (gBlocks, IDT). Genes were cloned from *E. coli* MG1655, *Vibrio fischeri* MJ1, *Rahnella aquatilis* ATCC 33071, *Sodalis praecaptivus* HS1, *Shigella flexneri* 2a, *Salmonella enterica enterica Typhimurium* ATCC 13311, or *Klebsiella pneumoniae* MGH 78578. The products were digested with EcoRI and HindIII and ligated into similarly digested pBAD24 or with NheI and KpnI and ligated into similarly digest pKT25c and pUT18c to generate N-terminally tagged T25 or T18 fusion proteins (43,69).

The Δ*mntP::mneA* replacement strain was generated by cloning the KanR cassette from pKD4 next to the *mneA* gene in pJS4B using the HiFi DNA Assembly cloning kit (NEB) (41). The *mneA*:KanR construct was amplified by PCR and moved into the chromosome cleanly replacing the *mntP* ORF via mini-λ Red recombineering (67).

Epitope-tagged constructs using the GST tag or the 3X FLAG tag were generated using the HiFi DNA Assembly cloning kit (NEB). MntS was cloned into pGEX-6-1 and the GST-MntS fusion was subcloned into pKT25c such that the resulting plasmid lacked the T25 region, or into pBAD24. Epitope tags were inserted into loop 3 of MntP between Glu95 and Asp96. The resulting MntP-3XFLAG construct was functional. The MntP-T18 and MntP-T25 constructs were less active than the untagged MntP, but could still partially rescue the manganese sensitivity defect of the *ΔmntP* strain.

### Metal Sensitivity Assays

The *ΔmntS* strain (LSW124) bearing the pBAD24 empty vector or indicated plasmids was grown overnight at 37 °C in LB-ampicillin, diluted 1:2000 into LB-ampicillin, and grown 2.5 hours to an OD600 of ∼ 0.25. Arabinose was added to induce gene expression and cells were grown for 30 min. Aliquots of 3 μL of 10-fold serial dilutions were spotted onto LB plates with ampicillin and arabinose containing the indicated concentrations of MnCl2, Zn(Ac)2, NiCl2, CoCl2, CuCl2, or Fe(NH4)2(SO4)2. Each spot represents an order of magnitude effect. In cells without a vector (Figure S1), no ampicillin was present. In cells with two-hybrid vectors (Figure S2), IPTG was also added to induce gene expression and kanamycin was used for cells containing the pKT25c vector.

### Bacterial Two-Hybrid Assays

Pairs of plasmids to be tested were fused to the two catalytic domains of T18 and T25 adenylate cyclase. Plasmids were co-transformed into *ΔcyaΔmntS* (LSW294) or *ΔcyaΔmntSΔmntP::mneA* (LSW304) strains and plated on LB-ampicillin-kanamycin at 30 °C. For each culture, three colonies were picked into LB-ampicillin-kanamycin supplemented with 0.5 mM IPTG and indicated concentrations of MnCl2, Zn(Ac)2, NiCl2, CoCl2, CuCl2, or Fe(NH4)2(SO4)2 and grown overnight at 30 °C. Beta-galactosidase assays were carried out as in (43,70). Briefly, the OD600 of the cultures was measured. Cells were permeabilized in Z-buffer containing BME, SDS, and CHCl3. The LacZ enzyme assay was initiated by addition of 100 μL of 8 mg/mL ONPG and terminated by 0.5 mL Na2CO3 and product formation was monitored using Abs420.

### Protein purifications

For MntS-MntP co-purifications, Δ*mntS* (LSW124) cells with pBAD24-MntP-3XFLAG or the empty vector pBAD24 and pKT25c-GST-MntS or the empty vector pKT25c were grown in LB to mid-log phase at 37 °C. GST-MntS expression was induced with 0.5 mM IPTG and MntS-3XFLAG expression was induced with 0.2% arabinose for 1-2 hours. Cells were harvested by centrifugation and lysed using B-PER detergent reagent (ThermoFisher) in Lysis Buffer (20 mM Tris pH 7.5, 50 mM NaCl, 5% glycerol) with 250 μg/mL lysozyme, 10 μg/mL DNaseI, 5 mM MgSO4, and 1 tablet per 40 mL EDTA-free protease inhibitors (Pierce). Insoluble material was removed by centrifugation at 20,000 xg for 20 min at 4 °C. Cleared lysate was applied to glutathione-sepharose (BioVision) or anti-FLAG M2 resin (Sigma) using 100 μL resin per 5 mL lysate and incubated 18 hours at 4 °C. Columns were washed with Lysis Buffer containing 2 mM DDM. Proteins were eluted using 10 mM glutathione or 0.5 mg/mL 3XFLAG peptide (Sigma) in Lysis Buffer with 2 mM DDM.

### Western Blotting

For Western blot analysis, samples in 1× SDS loading buffer with 100 mM dithiothreitol were not heated to avoid MntP aggregation. Proteins were separated on 14% Tris-Glycine gels and transferred to nitrocellulose membranes (Bio-Rad). Membranes were blocked with 3% milk in Tris-buffered saline with Tween-20 (TBS-T) and probed with 1:1000 anti-MntS (rabbit polyclonal) followed by 1:10,000 goat anti-rabbit-AzureSpectra800 (Azure) or 1:1000 anti-FLAG M2-HRP antibody (Sigma-Aldrich) in 3% milk-TBS-T. Signals were visualized using chemiluminescence (Bio-Rad) or emission at 832 nm on an Azure500 imager.

### Sequence Analysis

*Bona fide* MntS and SitA superfamily were obtained from the National Center for Biotechnology Information (NCBI) Genbank database. Sequence similarity searches were performed using the PSI-BLAST programs against the NCBI non-redundant (nr) database or the same database clustered down to 50% sequence identity using the MMseqs program with a profile-inclusion threshold was set at an e-value of 0.01 (71,72). Profile-profile searches were performed with the HHpred program (73). Multiple sequence alignments (MSAs) were constructed using the FAMSA and MAFFT programs (74,75). Sequence logos were constructed using these alignments with the ggseqlogo library for the R language (76). Signal peptides were predicted using a deep neural network as implemented in SignalP 6 (77).

### Structure Analysis

The JPred program was used to predict secondary structures using MSAs (see above). PDB coordinates of structures were retrieved from the Protein Data Bank. Structures were rendered, compared, and superimposed using the Mol* program (78). Structural models were generated using the RoseTTAfold and Alphafold2 programs (79,80). Multiple alignments of related sequences (>30% similarity) were used to initiate HHpred searches for the step of identifying templates to be used by the neural networks deployed by these programs.

### Comparative Genomics and Phylogenetic Analysis

Clustering of protein sequences and the subsequent assignment of sequences to distinct families was performed by the MMSEQS program adjusting the length of aligned regions and bit-score density threshold empirically. Divergent sequences or small clusters were merged with larger clusters if other lines of evidence, including shared sequence motifs, shared structural synapomorphies, reciprocal BLAST search results, and/or shared genome context associations, supported their inclusion. Phylogenetic analysis was performed using the maximum-likelihood method with the WAG or Blosum62 models with the IQTree and FastTree program (81,82). For example, a tree of the SitA metal-binding protein was generated with the WAG substitution matrix using the command FastTree -close 0.618 -cat 15 -refresh 0.7 -pseudo -wag. The FigTree program (http://tree.bio.ed.ac.uk/software/figtree/) was used to render phylogenetic trees. Gene neighborhoods were extracted through custom PERL scripts from genomes retrieved from the NCBI Genome database. These were then clustered using MMSEQS and filtered using a neighborhood distance cutoffs and phyletic patterns to identify conserved gene neighborhoods.

## Acknowledgements

We thank M. Hemm, M. Wu Orr, R. Zeinert, and G. Storz for helpful discussions. This work is supported by Grant R15GM137249 from the NIH (to LSW), Grant 23595 from the Research Corporation for Science Advancement (to LSW), and the Intramural Research Program of the National Library of Medicine (VA and LA).

---

**Supplemental Figure 1:**
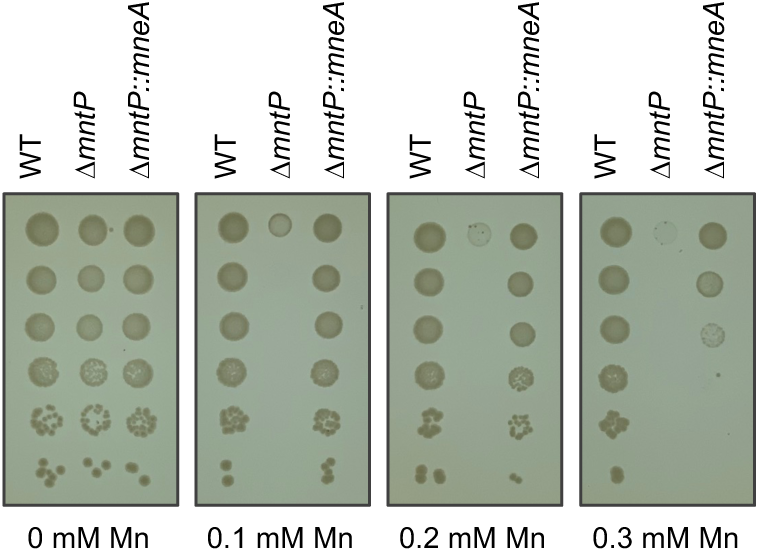
The heterologous MneA Mn exporter can rescue growth of a Δ*mntP* strain under supplemental Mn conditions. Mn sensitivity assay showing that replacement of the endogenous *mntP* gene encoding the *E. coli* Mn exporter with a heterologous *mneA* gene encoding an unrelated *Vibrio fischerii* Mn exporter restores growth of the *ΔmntP* strain in the presence of Mn. Ten-fold serial dilutions of mid-exponential-phase cultures of WT, Δ*mntP* cells, or *ΔmntP::mneA* cells were spotted onto LB plates containing the indicated amounts of MnCl_2_. Results are representative of at least three independent replicates.

**Supplemental Figure 2:**
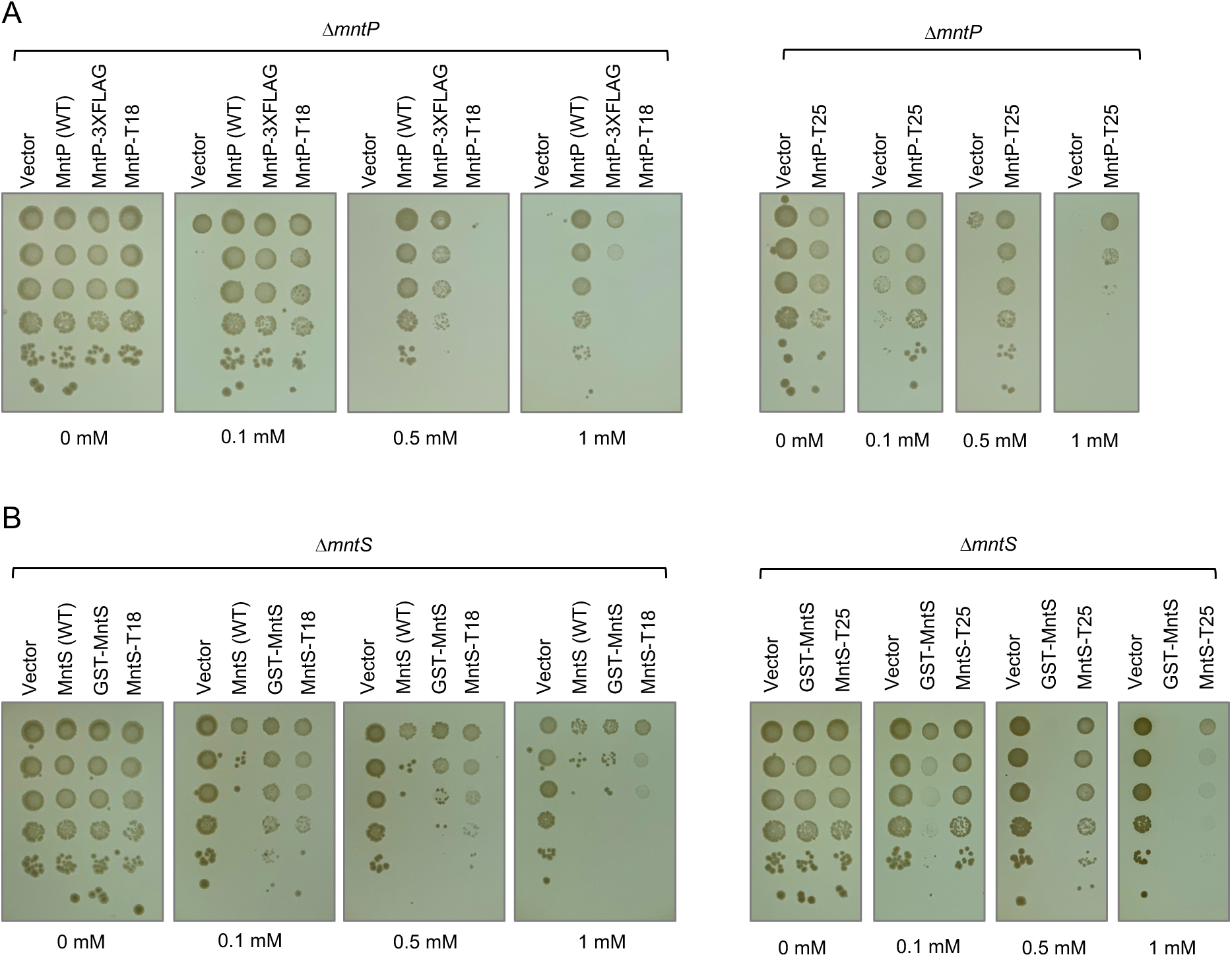
Epitope-tagged MntP and MntS constructs retain functionality. **A)** Functionality of epitope-tagged versions of *mntS* and *mntP* was assessed using a Mn sensitivity assay. Ten-fold serial dilutions of mid-exponential-phase cultures of Δ*mntP* cells bearing empty vectors (pBAD24 or pKT25c), pBAD24 with MntP, MntP-3XFLAG, and the two-hybrid vectors (pUT18c or pKT25c) with MntP-T18 or MntP-T25 respectively were spotted onto LB-ampicillin or LB-kanamycin plates containing the indicated amounts of MnCl_2_. Results are representative of at least three independent replicates. **B)** The Mn sensitivity assay was carried out as above with Δ*mntS* cells bearing empty vectors (pBAD24 or pKT25c), pBAD24 with MntS or GST-MntS, and the two-hybrid vectors (pUT18c or pKT25c) with T18-MntS or T25-MntS, as well as the pKT25c vector lacking the T25 domain and instead expressing GST-MntS.

**Supplemental Figure 3:**
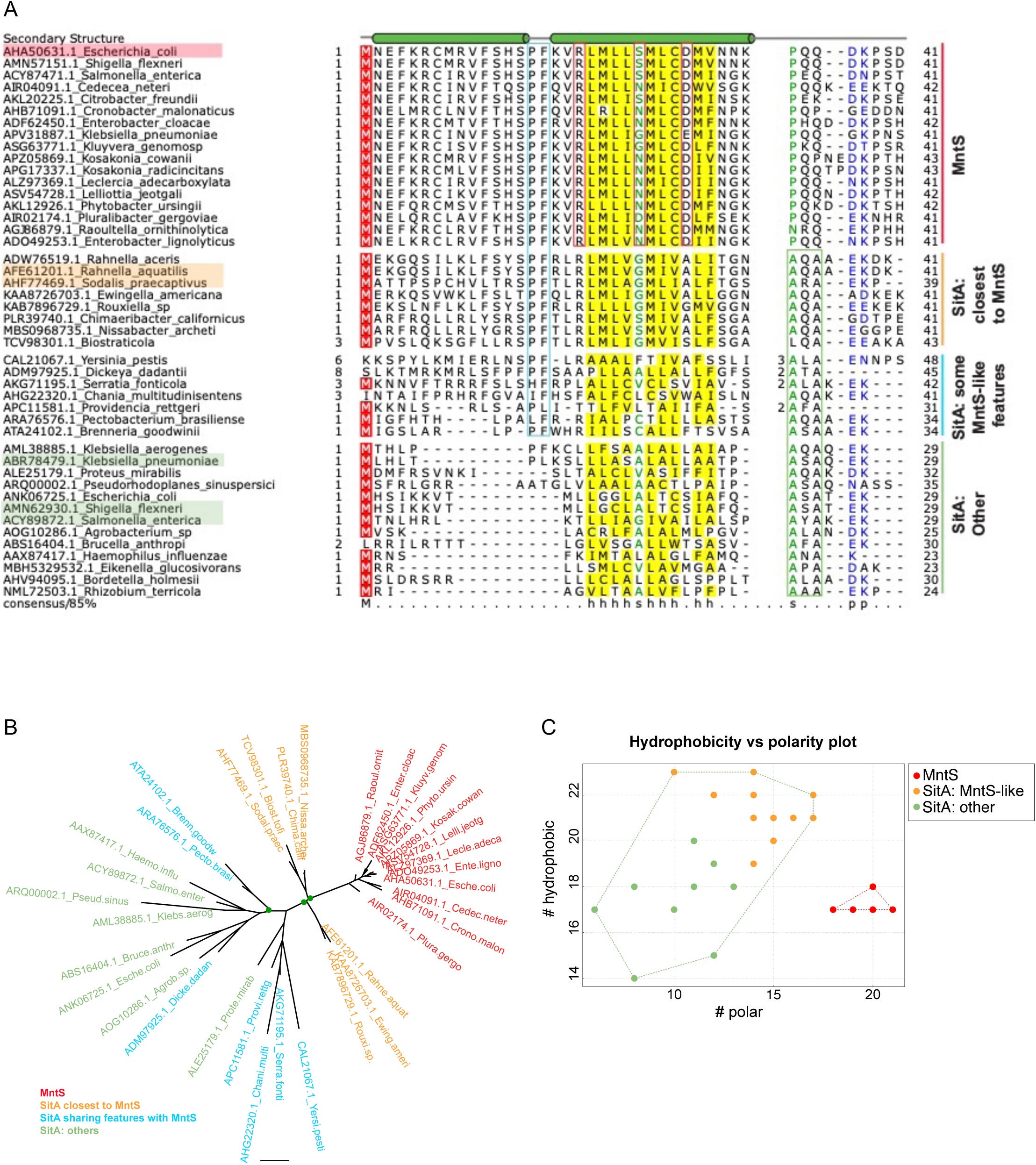
Multiple sequence alignment, phylogeny, and hydrophobicity plot of MntS and SitA SPs. **A)** Multiple sequence alignment of the MntS and SitA SP grouped by relatedness to MntS (shown on right). A set of 17 MntS and 28 SitA SPs were excerpted from a comprehensive collection of MntS and SitA proteins (see legend to Supplemental Figure 4). The accessions and organism names are given. The secondary structure is shown on the top where the green cylinder represents a helix. The residues are colored by the 85% consensus shown in the last line. The amino acid classes in the consensus are colored as follows: polar (p: KRHEDQNST) in blue; hydrophobic (h: ALICVMYFW) with yellow background; small (s: ACDGNPSTV) in green; and the conserved ‘M’ with a red background. The limits of the domains are indicated by the amino acid numbers on each side, and inserts in some sequences are shown as numbers. The PF marking the end of an inserted helix is in a blue box; polar residues conserved in the hydrophobic core of MntS are in a red box; AXA-motif is in a green box. The *E. coli* MntS sequence is indicated in red, the highly related MntS-like SitA SP sequences cloned from *Rahnella* and *Sodalis* are highlighted in orange, and distant SP sequences cloned from *Shigella*, *Salmonella*, and *Klebsiella* are highlighted in green. **B)** A phylogenetic tree of constructed using the non-redundant set of sequences from the above alignment showing the nesting of the MntS within a clade of SitA SPs that share specific sequence motifs with them. The tree was generated using IQtree with the Blosum62 matrix as the alignment is very short. Key branches with bootstrap values greater than 80% are shown with green dots. **C)** The total number of hydrophobic positions in representative MntS or SitA SPs is plotted against the total number of polar positions in the same. Note the systematic shift in MntS toward a more polar state as opposed to the SPs, which likely indicates selection for greater solubility and potential cytoplasmic function.

**Supplemental Figure 4:**
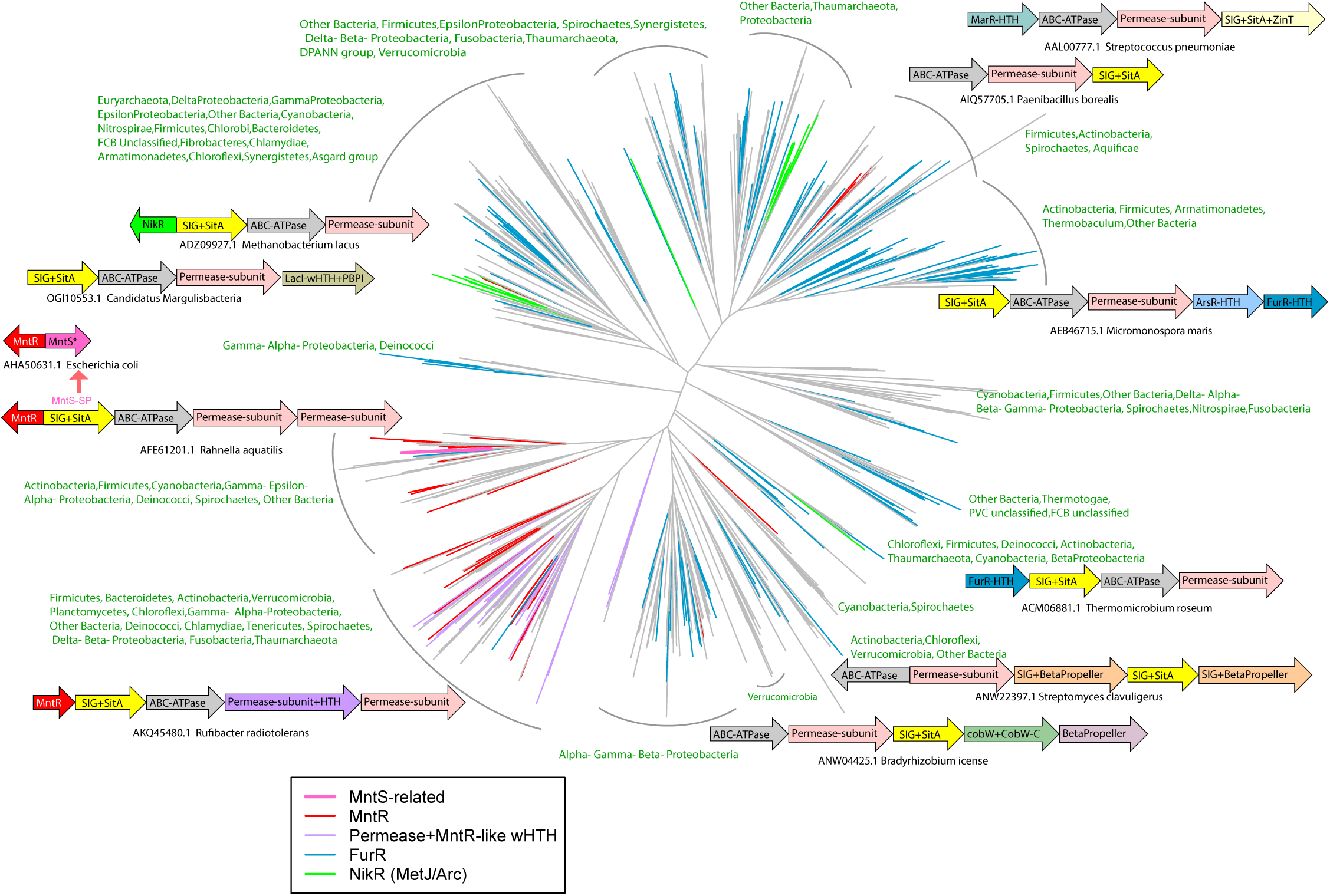
Phylogenetic tree of SitA SP showing a detailed distribution of gene-neighborhood themes and phyletic patterns. A phylogenetic tree of SitA with representative operons was generated as in Figure 3B. A comprehensive set of MntS and SitA proteins were collected from a curated database of 4210 complete prokaryotic genomes and 2983 metagenomes augmented with addition sequences whole-genome shotgun sequences from the NCBI non-redundant database. This collection included 3736 SitA proteins and 103 MntS proteins. A 50% clustering by MMseqs followed by manual selection of unfragmented and phyletically representative sequences gave 506 that were used for the above tree. An expanded version of phyletic patterns (relative to Figure 3B) of the major branches are shown in green and the tree colored as described in the legend to Figure 3. Abbreviations are as in Figure 3B with the following additional ones: SIG – SitA signal peptide region; LacI, MarR, ArsR – different families of wHTH DNA binding domains prototyped by the eponymous TFs; PBPI – Periplasmic Binding Protein-I-type ligand-sensor domain; CobW – CobW/HypB/UreG-like GTPase domain; CobW-C: C-terminal domain required for metal-insertion by these GTPases; ZinT – ZinT heavy metal-binding domain; β-propeller – WD40 repeat protein.

**Supplemental Figure 5:**
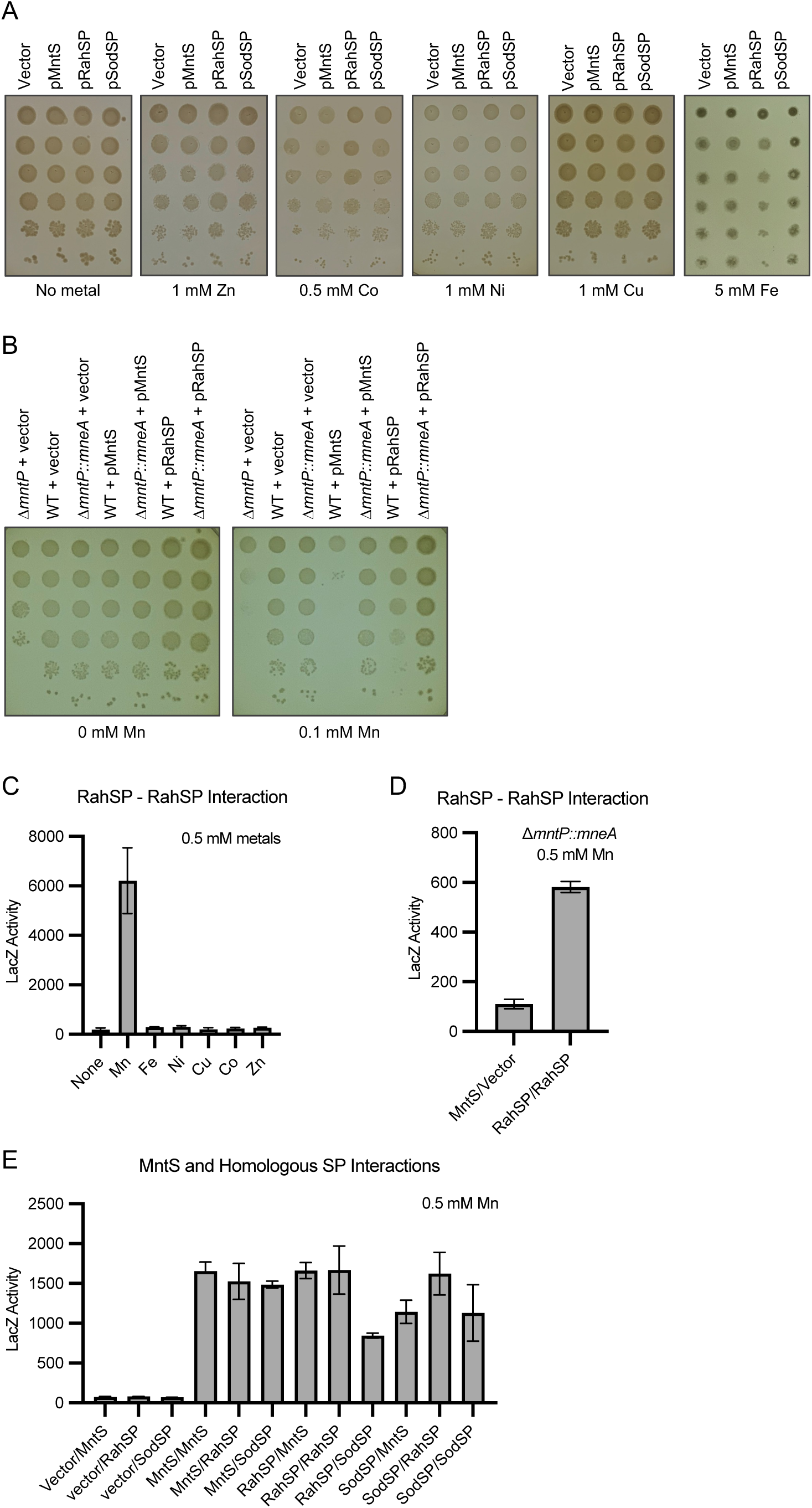
MntS-like SP regions of SitA act similarly to MntS. **A)** Metal sensitivity assays showing that the *Rahnella* and *Sodalis* SP regions only cause a growth defect in the presence of Mn and not other metals. Ten-fold serial dilutions of mid-exponential-phase cultures of Δ*mntS* cells bearing the pBAD24 empty vector or pBAD24 expressing MntS, RahSP, or SodSP were spotted onto LB plates containing the indicated amounts of metals. Results are representative of at least three independent replicates. **B)** Mn sensitivity assay showing that the *Rahnella* SP region can only cause a Mn-dependent growth defect in cells producing the MntP protein. Overexpression of MntS or the *Rahnella* SP region causes a growth defect in the presence of Mn in the WT strain background, but the Mn sensitivity for both MntS and the *Rahnella* SP region is lost in the Δ*mntP::mneA* strain which lacks the MntP exporter. Ten-fold serial dilutions of mid-exponential-phase cultures of WT or Δ*mntP::mneA* cells bearing the pBAD24 empty vector, pBAD24-MntS, or pBAD24-RahSP were spotted onto LB plates containing the indicated amounts of MnCl_2_. Results are representative of at least three independent replicates. **C)** Two hybrid assay showing that *Rahnella* SP region only self-interacts in the presence of Mn and not other metals in Δ*cya* Δ*mntS* cells. For all two-hybrid assays (C-E), cells containing two plasmids with fusions of MntS or indicated SP regions to the T18 and T25 fragments of adenylate cyclase were grown overnight in LB media with 0.5 mM of indicated metals. β-galactosidase assays were performed to quantify the LacZ activity representing the protein-protein interaction. Results are given in Miller units as the mean ± SD of three independent samples and are representative at least three independent experiments. **S)** Two-hybrid assay showing that the *Rahnella* SP region still interacts with itself in the Δ*cya* Δ*mntS ΔmntP::mneA* strain which lacks the MntP exporter. **E)** Two hybrid assay showing that MntS and homologous SP regions from SitA proteins (*Rahnella* and *Sodalis*) can heterodimerize with each other in all combinations in Δ*cya* Δ*mntS* cells.

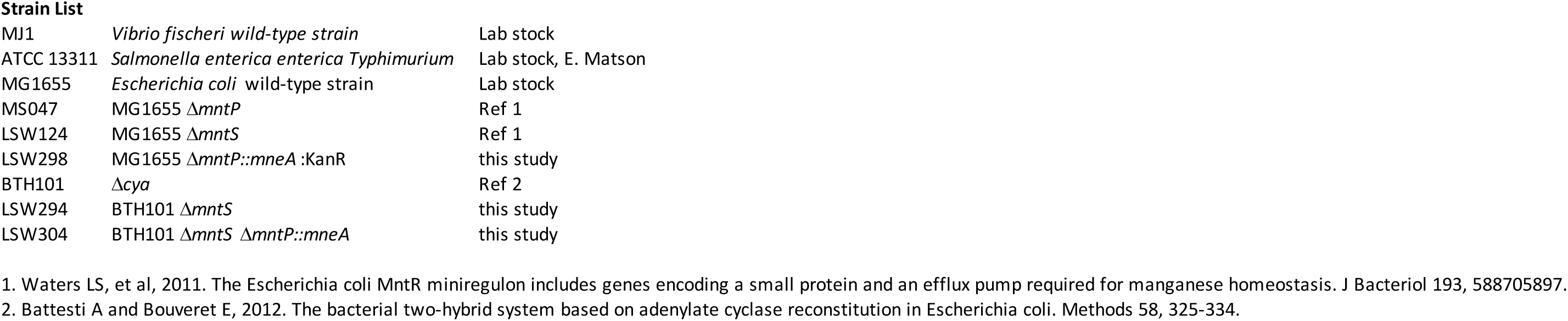
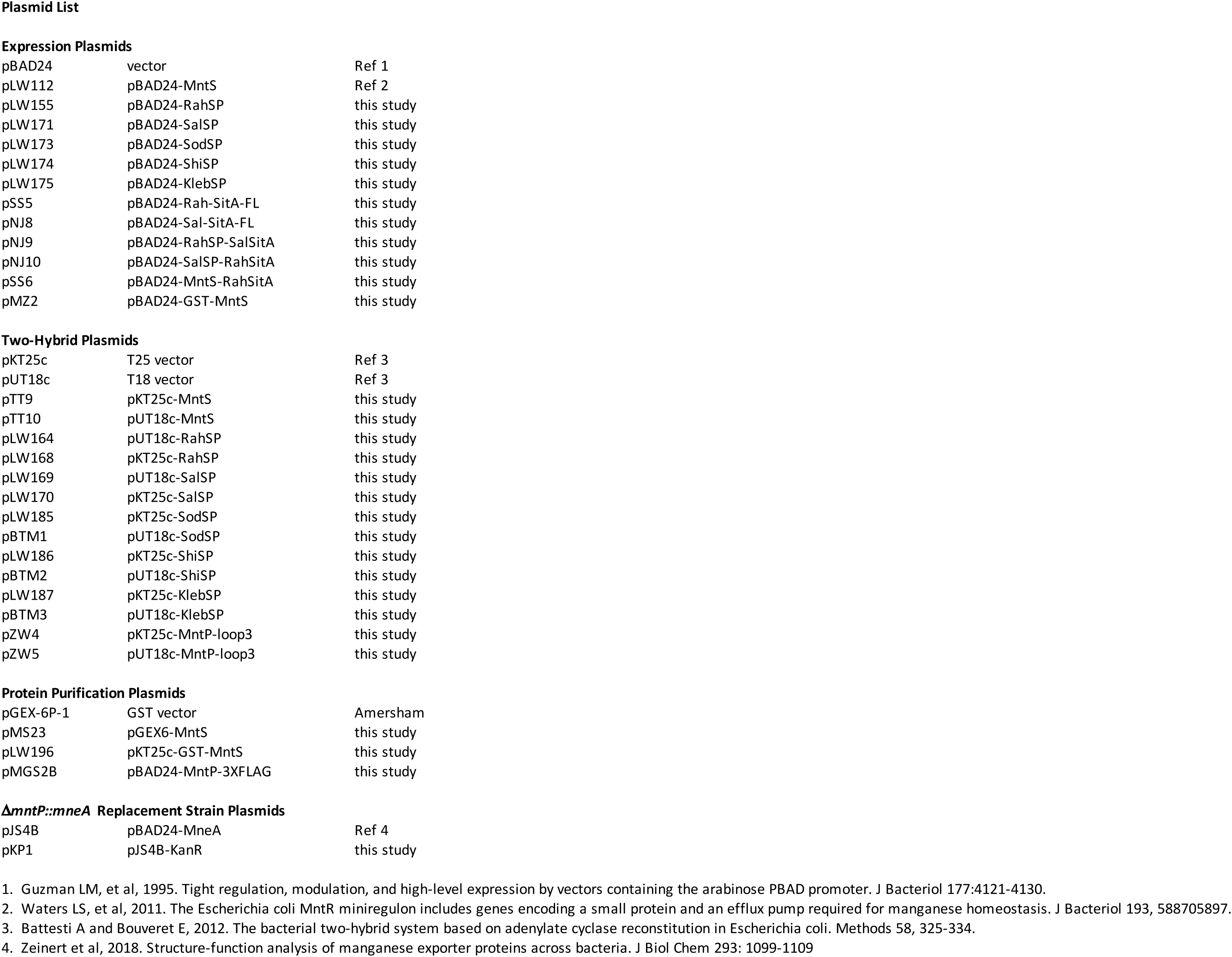
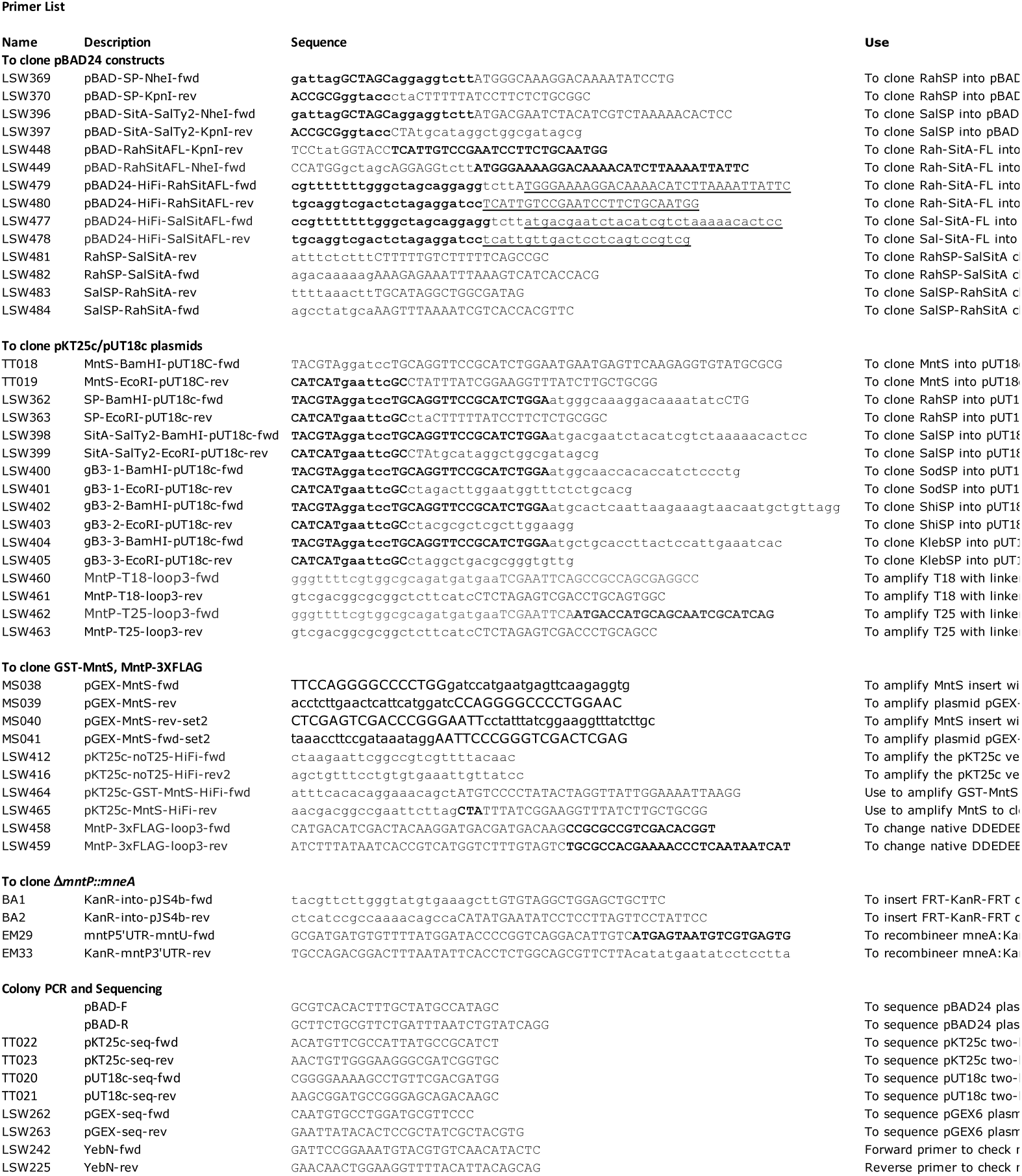
Supplemental Tables for Wright *et al*, The small protein MntS evolved from a signal peptide sequence and acquired a novel function in Mn homeostasiss.

